# NK cells prevent the formation of teratomas derived from human induced pluripotent stem cells

**DOI:** 10.1101/714170

**Authors:** Basma Benabdallah, Cynthia Désaulniers-Langevin, Chloé Colas, Yuanyi Li, Jean V. Guimont, Elie Haddad, Christian Beauséjour

## Abstract

The safe utilization of induced pluripotent stem cell-derivatives in clinic is tributary to the complete elimination of the risk of forming teratomas after transplantation. The extent by which such a risk exists in immune competent hosts is mostly unknown. Here, using humanized mice reconstituted with fetal hematopoietic stem cells and autologous thymus tissue (Hu-BLT) or following the adoptive transfer of peripheral blood mononuclear cells (PBMCs) (Hu-AT), we evaluated the capacity of immune cells to prevent or eliminate teratomas derived from human induced pluripotent stem cells (hiPSCs). Our results showed that the injection of hiPSCs failed to form teratomas in Hu-AT mice reconstituted with allogeneic or autologous PBMCs or purified NK cells alone. However, teratomas were observed in Hu-AT mice reconstituted with autologous PBMCs depleted from NK cells. In line with these results, Hu-BLT which do not have functional NK cells could not prevent the growth of autologous teratomas. Finally, we found that established teratomas were not targeted by NK cells and instead were efficiently rejected by allogeneic but not autologous T cells in Hu-AT mice. Overall, our findings suggest that autologous hiPSC-derived therapies are unlikely to form teratomas in the presence of NK cells.

## INTRODUCTION

The potential of regenerative medicine is greatly enhanced by the development of induced pluripotent stem cells (iPSCs). Yet, the risk of forming teratomas using iPSC-derived cell grafts could compromise their clinical use. Indeed, the injection of only a few hundred human embryonic cells was sufficient to form a teratoma in immunodeficient mice [1]. To prevent the formation of teratomas, small molecules able to selectively and efficiently kill pluripotent cell by inhibiting antiapoptotic factors were developed [2; 3]. However, the need for pluripotent stem cells to evade immune responses may be required for the growth of teratomas *in vivo*. Pioneer work by Dressel and colleagues showed that mouse iPSCs can occasionally give rise to teratomas in autologous/syngeneic mice in absence of the activation of the host natural killer (NK) cells [4; 5]. Likewise, human iPSCs were shown to be the target of human NK cells *in vitro* [6; 7]. Yet, the *in vivo* contribution of the innate immunity, particularly the role of NK cells in the tumorigenic potential of hiPSCs, remains unknown. Here, using two different models of humanized mice, we show that teratoma formation by human iPSCs is abolished only in the presence of NK cells and that this NK-specific cytotoxicity is lost upon the differentiation of hiPSCs.

## EXPERIMENTAL PROCEDURES

### Humanized mice

NOD/SCID/IL2Rγ null (NSG) mice were obtained from the Jackson Laboratory (Bar Harbor, ME) and housed in the animal care facility at the CHU Sainte-Justine Research Center under pathogen-free conditions in sterile ventilated racks. All *in vivo* manipulations were previously approved by the institutional committee for good laboratory practices for animal research (protocol #579). Bone-liver-thymus humanized mice (Hu-BLT) were generated as previously described [8]. Briefly, six week-old NSG mice were first irradiated with 2 Gy total body irradiation (1 Gy/min using a Faxitron CP◻160) and implanted with small pieces (1-2 mm^3^) of human fetal thymus under the renal capsule followed by the intravenous delivery of 1×10^7^ CD34^+^ hematopoietic stem cells isolated from autologous fetal liver. Fetal tissues were obtained from consented healthy donors after surgical abortion at around week 20 of pregnancy. Human immune cell engraftment in humanized mice was monitored in peripheral blood until 13 weeks post-reconstitution. Leukocytes were labeled with conjugated antibodies for human PerCP-Cy5.5-CD45, APC-CD3, PE-CD19, and FITC-CD4 (all from BD Biosciences) and analyzed by flow cytometry (BD FACSCANTO II, BD Biosciences). For adoptive transfer experiments (Hu-AT), human adult blood was collected and immune cells purified by Ficoll (GE healthcare). Mice were injected intravenously with 1×10^7^ freshly isolated PBMCs or NK-depleted PBMCs obtained from the negative fraction of a positive selection (CD56+) kit (STEMCELL Technologies). Alternatively, mice were injected with 5-15×10^5^ NK cells purified using the NK cell enrichment negative selection kit (STEMCELL Technologies).

### Generation and characterization of hiPSCs

Fibroblasts isolated either from human fetal liver tissues or human adult skin after collagenase dissociation were reprogrammed into iPSCs with the integration-free based Sendai virus (Cytotune 2.0 kit from Life Technologies). Fibroblasts were used at low population doubling (less than 5) to insure high efficiency of reprogramming. Emerging hiPSC colonies were manually picked and cultured under feeder-free conditions in Essential 8 medium on Geltrex-coated dishes (Life Technologies). hiPSC clones were passaged at least 15 times to increase stable pluripotency. hiPSC generation and characterization were done in the iPSC – cell reprogramming core facility of CHU Sainte-Justine. hiPSC colonies were stained with antibodies for anti-human SSEA-4, Sox2, OCT4 and TRA1-60 overnight at 4°C using the pluripotent Stem Cell 4-Marker Immunocytochemistry Kit (Life Technologies), followed by incubation with an ALEXA secondary antibodies for 30 minutes at room temperature. Nuclei were counterstained with DAPI. Karyotypes were produced by G-banding and analyzed by the CHU Ste-Justine cytogenetic department.

### T cell activation and proliferation assays

Human PBMCs were freshly isolated from healthy donors peripheral blood using Ficoll. Effector cells were then co-cultured with either autologous or allogeneic iPSCs at 1:2 ratio during three days for T cell activation or 5 days for T cell proliferation at 37°C. T cell activation was measured with a PE-conjugated anti-hCD69 on CD3^+^ gated viable cells by flow cytometry (BD LSRFortessa, BD Biosciences). For T cell proliferation, PBMCs were first stained using the CellTrace CFSE kit to monitor distinct generations of viable proliferating CD3^+^ T cells by dye dilution (Invitrogen) before being co-cultured with iPSCs. Effector cells without stimulation were used as a negative control and an anti-CD3 (OKT3) antibody were used as positive control. 7-AAD (BD Biosciences) was used to exclude dead cells.

### NK cell degranulation and cytotoxicity assays

Freshly purified NK cells (as described above) were incubated with or without target cells at the indicated ratios. K562 cells were used as a positive control in all experiments. For the NK cell degranulation assay, effector and target cells were co-cultured at 1:2 ratio in the presence of FITC-conjugated anti-human CD107a/b for one hour at 37°C, then 2μl/ml of monensin (BD Biosciences) was added to the cell mixture for an additional three hours of incubation. For cytotoxicity assay, effector and PKH26-stained target cells were mixed at 1:1 or 5:1 ratio and incubated for four hours at 37°C. At the end of the incubation, degranulation was quantified by flow cytometry (BD LSRFortessa, BD Biosciences) after gating on CD3−/CD56+/CD107+ viable cells and the extent of cytotoxicity was determined by the relative number of live target cells labelled with PKH26 only and dead cells labelled with both PKH26 (Sigma-Aldrich) and 7-AAD (BD Biosciences).

### Teratoma formation

Approximately 1×10^6^ hiPSCs were first detached in cell clumps using the gentle cell dissociation reagent (STEMCELL Technologies) and resuspended in cold PBS containing 50% of Geltrex and kept on ice. Cell mixture was then injected unilaterally under the renal capsule of mice. Eight weeks later, teratomas were dissected out and placed in OCT. Cryosections were stained with hematoxylin eosin for histological analysis, or immunostained with antibodies against human CD4, CD8, NKp46 (all from Biolegend) and TUNEL (Roche).

### Statistical Analysis

GraphPad Prism 8 software was used for statistical analysis; ρ values on multiple comparisons were calculated using Student’s t-Tests

## RESULTS

### NK cells prevent the formation of hiPSC-derived teratomas in Hu-AT mice

To investigate if immune cells could protect against the formation of hiPSC-derived teratomas, we first generated hiPSC clones from skin fibroblasts or PBMCs collected from three donors using an integration-free (Sendai virus) approach. We confirmed that these clones had normal karyotypes, expressed the classical markers of pluripotency and were able of forming the three germ layers in NSG immunodeficient mice (Figure S1A and S1B and data not shown). We then injected various hiPSC clones under the renal capsule of NSG mice and determined if the adoptive transfer of 1×10^7^ PBMCs could prevent the formation of teratomas (Figure 1A). Our results show that no teratomas were found after the adoptive transfer of autologous or allogeneic PBMCs (Figure 1B). To better define the role of T and NK cells in this process, we repeated the experiment using either NK-depleted PBMCs or purified NK cells (Figure S2). We found that hiPSCs efficiently formed teratomas in the presence of autologous PBMCs depleted from NK cells. This was in contrast to when mice were injected with allogeneic NK-depleted PBMCs, in which case teratomas were rejected. This can be explained if in the absence of NK cells, hiPSCs were allowed to grow into teratomas and that differentiated cells were then rejected by allogeneic T cells. Consistent with that, *in vitro* assays showed that hiPSCs were the target of NK cells but were unable to activate T cells (Figures 2A-D). These results are also supported by our observation that hiPSC-derived myogenic cells can induce activation of allogeneic but not of autologous T cells in humanized mice (Benabdallah et al, 2019 submitted). This suggests that hiPSCs are prone to NK but not to T cell killing, an effect that could be mediated by the presence of NK activating receptors and low expression of HLA-I and costimulatory molecules at the surface of hiPSCs (Figure S1C). Intriguingly, we observed that mice injected with NK-depleted PBMCs had significantly larger teratomas compared to those grown in immunodeficient mice (Figure 1C). We speculate that their enhanced growth was fueled by human cytokines secreted by autologous PBMCs. Importantly, our results also show that the injection of purified NK cells (5-15×10^5^) was able to prevent the formation of hiPSC-derived teratomas in both autologous and allogeneic settings (Figure 1B). This confirms that NK cells, but not autologous T cells, are efficient in preventing the development of hiPSC-derived teratomas.

**Figure 1.**
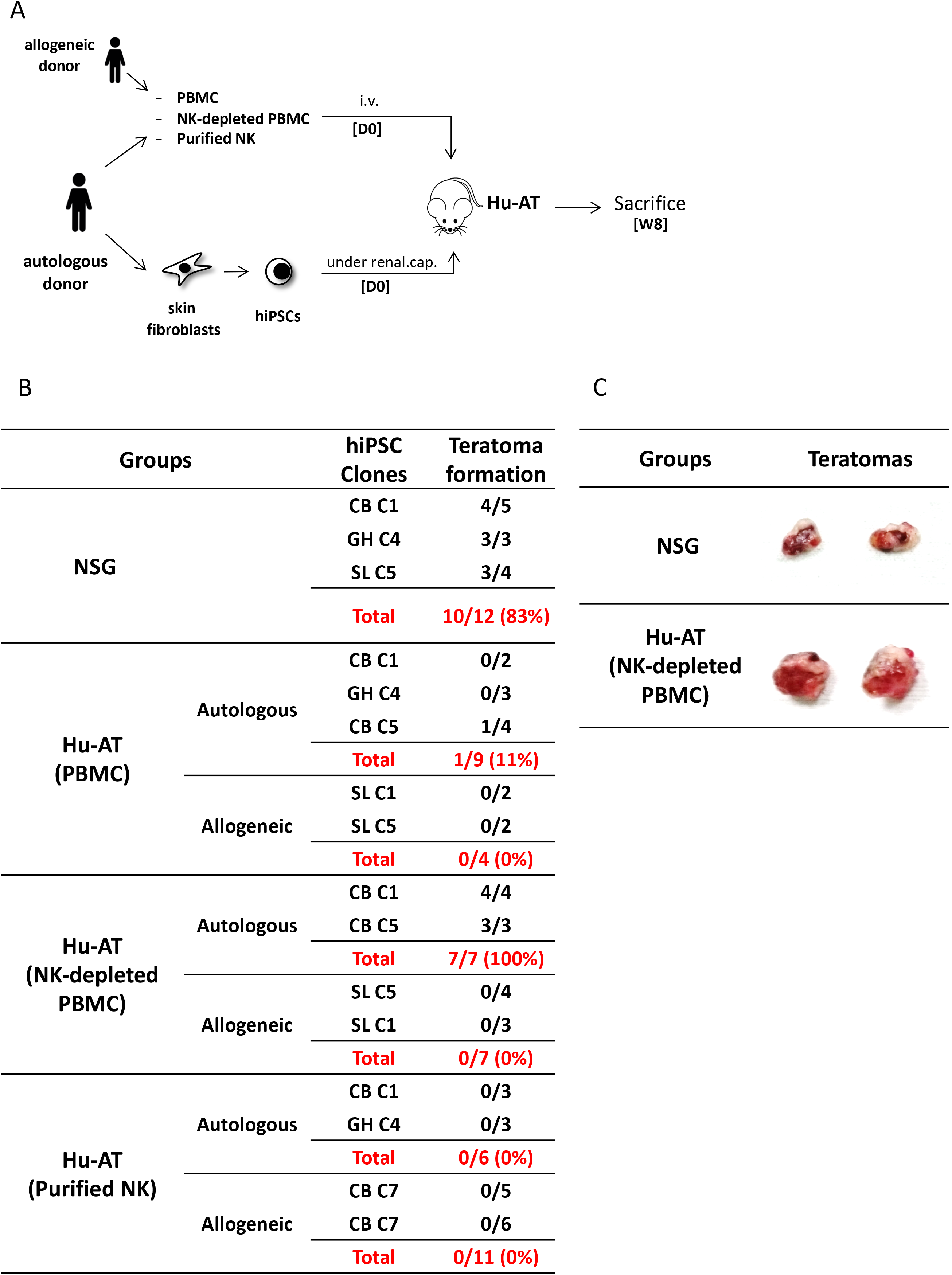
Injection of NK cells is sufficient to prevent the formation of hiPSC-derived teratomas in Hu-AT mice. **(A)** Experimental design and timeline of the injection of hiPSCs in Hu-AT mice. PBMCs (1×10^7^), NK-depleted PBMCs (1×10^7^) or purified NK cells (5-15×10^5^) were injected intravenously at day 0 (DO) to generate Hu-AT mice. Approximately 1×10^6^ autologous or allogeneic hiPSCs were injected under the renal capsule on the same day (one injection per mouse). Teratoma formation was evaluated upon sacrifice 8 weeks after the injection of cells. **(B)** Incidence of teratoma formation in the renal capsule of Hu-AT mice. Shown is the proportion of teratomas derived from the indicated hiPSC clones. **(C)** Representative photos showing the increased size of teratomas isolated from NSG mice previously injected with NK-depleted autologous PBMCs compared to control NSG non-reconstituted mice.

**Figure 2.**
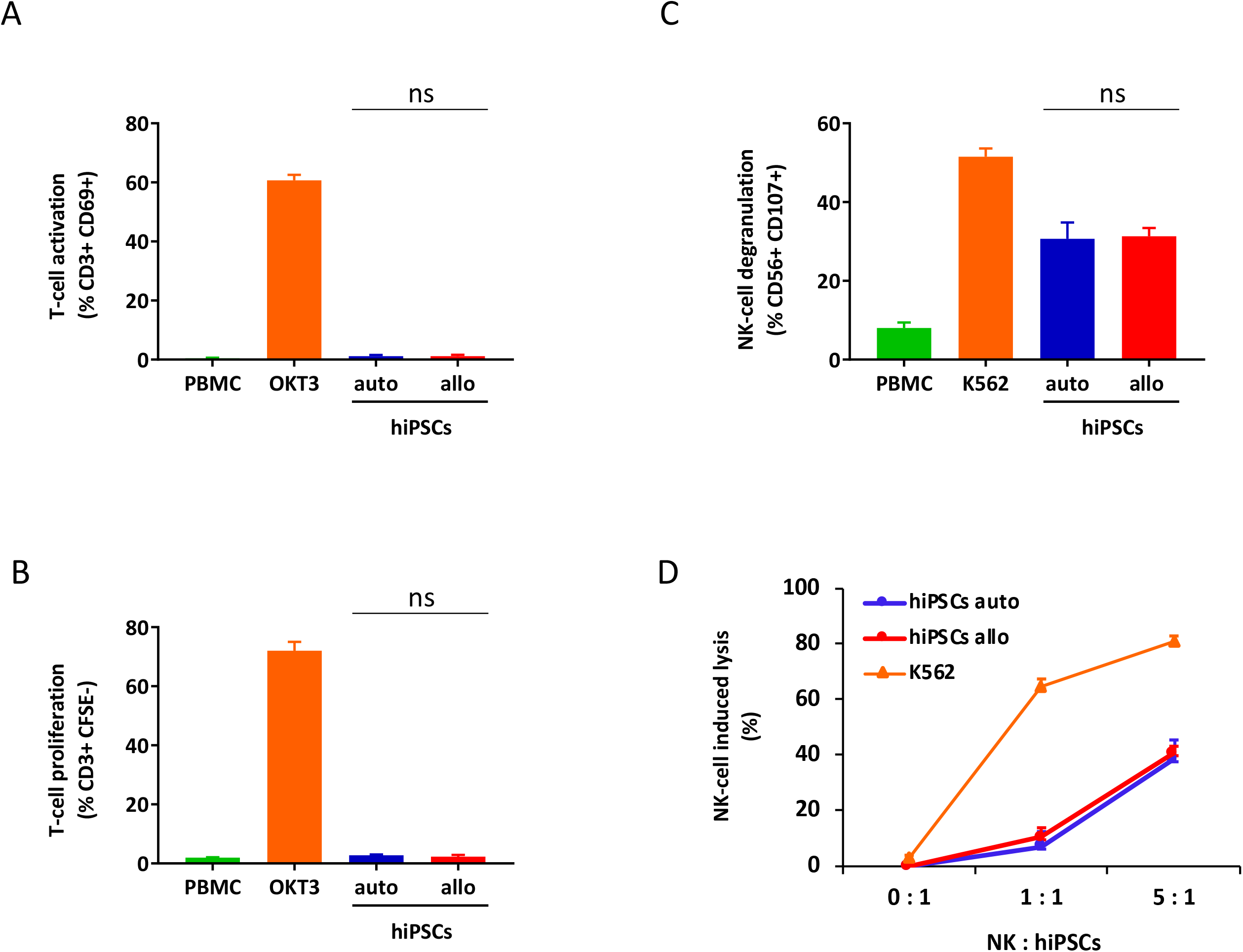
hiPSCs are the target of NK but not T cells *in vitro.* **(A)** Measures of T cell activation (as determined by gating for CD3^+^/CD69^+^ cells) after co-culture of hiPSCs with autologous or allogeneic PBMCs for 3 days. PBMCs alone were used as a negative control and PBMCs treated with OKT3 were used as a positive control. Shown is the mean ± SEM of 2 independent experiments done in triplicate using cells collected from 2 different donors. **(B)** Measures of T cell proliferation (as determined by gating for CD3^+^/CFSE^+^ cells) after co-culture of hiPSCs with autologous or allogeneic CFSE-stained PBMCs for 5 days. PBMCs alone were used as a negative control and PBMCs treated with OKT3 were used as a positive control. Shown is the mean ± SEM of 2 independent experiments done in triplicate using cells collected from 2 different donors. **(C)** hiPSCs induce degranulation of NK cells *in vitro*. Degranulation was determined by evaluating CD107 expression on gated CD3^−^/CD56^+^ NK cell populations after a 4 hour co-culture between freshly isolated PBMCs and autologous or allogeneic hiPSCs (ratio 1:2). PBMCs alone were used as a negative control and PBMCs co-cultured with K562 cells were used as a positive control. Shown is the mean ± SEM of 2 independent experiments done in triplicate using cells collected from 2 different donors. **(D)** Autologous and allogeneic hiPSCs are lysed by purified NK cells *in vitro*. Cell lysis was determined by flow cytometry with the absolute count of PKH26-stained hiPSCs after a 4 hour co-culture with NK cells purified by magnetic negative selection. NK cells alone were used as a negative control and NK cells co-cultured with K562 cells were used as a positive control. Shown is the mean ± SEM of 2 independent experiments done in triplicate using cells collected from 2 different donors.

### hiPSCs efficiently form teratomas in Hu-BLT mice lacking functional NK cells

To further investigate the role of NK cells in the development of hiPSC-derived teratomas, we then injected hiPSCs under the renal capsule of fully reconstituted Hu-BLT mice generated after the transplantation of fetal CD34 cells and thymus tissue fragments (Figure 3A). Hu-BLT mice are known to support reconstitution, maturation, and selection of T cells, but lack functional NK cells (Figure S3) [9]. Consistent with our findings in Hu-AT mice, our results show that the growth of autologous hiPSC-derived teratomas was not hampered in Hu-BLT mice (Figure 3B). Teratomas formed in Hu-BLT mice injected with autologous immune cells were found infiltrated with low level, non-randomly distributed T cells with no significant sign of apoptosis (Figure 3C), supporting the findings of Zhao and colleagues [10]. This is in contrast to when hiPSCs were injected in Hu-BLT mice reconstituted with allogeneic immune cells, in which the growth of teratomas was partially inhibited (50%) and where residual teratomas were massively infiltrated by T cells and undergoing high levels of apoptosis (Figure 3C).

**Figure 3.**
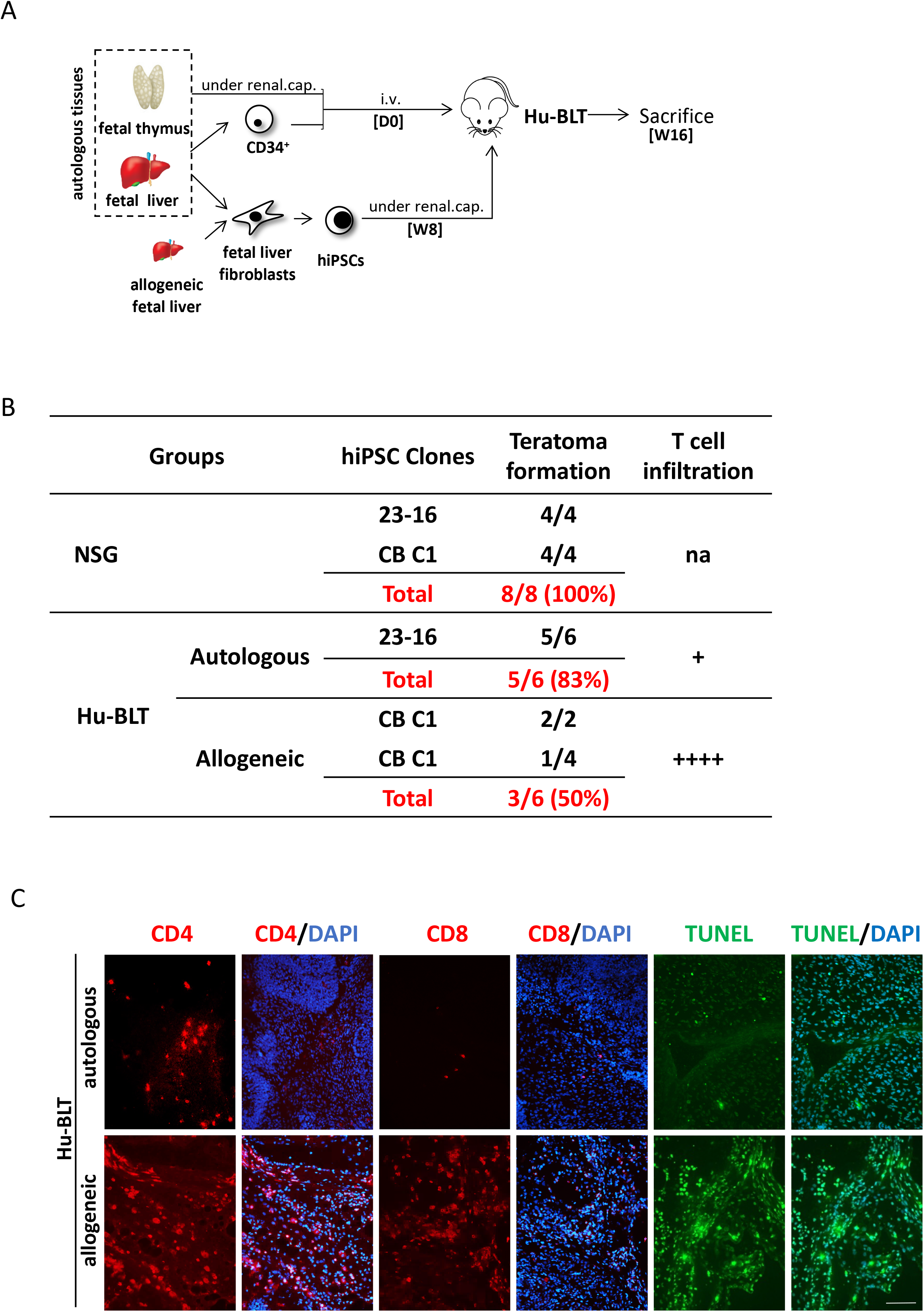
hiPSCs efficiently develop into teratomas in Hu-BLT mice lacking functional NK cells. **(A)** Experimental design and timeline of the injection of hiPSCs in Hu-BLT mice. Hu-BLT mice were generated by the injection of fetal tissues as previously described and immune reconsititution allowed for 8 weeks (W8) prior from the injection of approximatively 1×10^6^ hiPSCs under the renal capsule. hiPSCs were derived from fetal liver fibroblasts autologous or allogeneic to CD34 positive hematopoietic cells. Teratomas were allowed to grow for 8 weeks before mice were sacrificed at week 16 (W16). **(B)** Incidence of teratoma formation in the renal capsule of Hu-BLT mice. Shown is the proportion of teratomas derived from the indicated hiPSC clones and their relative infiltration with T cells. na = not applicable. **(C)** Representative photos showing the infiltration of CD4^+^ and CD8^+^ T cells (in red) and apoptosis (as determined by TUNEL in green) in sections of allogeneic and autologous teratomas collected from Hu-BLT mice. DAPI staining was performed to visualize nuclei (in blue). Showed are photos taken at 20X. Scale bar, 100 μm.

### NK cells fail to reject established teratomas

We then hypothesised that the protection against the formation of teratomas conferred by NK cells was tributary to the undifferentiated nature of iPSCs (i.e. low HLA-I expression) which favors the activation of NK cells. To test this hypothesis, we repeated the above-described experiment but this time we proceeded to the adoptive transfer of immune cells only six weeks after the injection of hiPSCs, a time at which teratomas had already started to grow and cells to differentiate (Figure 4A). While we found that allogeneic PBMCs or NK-depleted PBMCs were capable of rejecting teratomas, autologous PBMCs and purified NK cells were not (Figure 4B). Analysis of the teratomas upon sacrifice showed a massive T cell infiltration and apoptosis in the remaining allogeneic teratomas (Figure 4C). In comparison, only a few T cells were found in autologous teratomas. As expected, very few NK cells were found in growing teratomas, suggesting these cells lose their ability to recognize differentiated hiPSC-derived cells. This is supported by the fact that while hiPSCs expressed very low levels of HLA-I expression, these levels were found much higher within only a few hours after the cells were placed in presence of serum *in vitro* (Figure S4). Overall, these results indicate that NK cells are very effective at preventing the formation of a teratoma from hiPSCs, but lose this capacity against differentiated cells.

**Figure 4.**
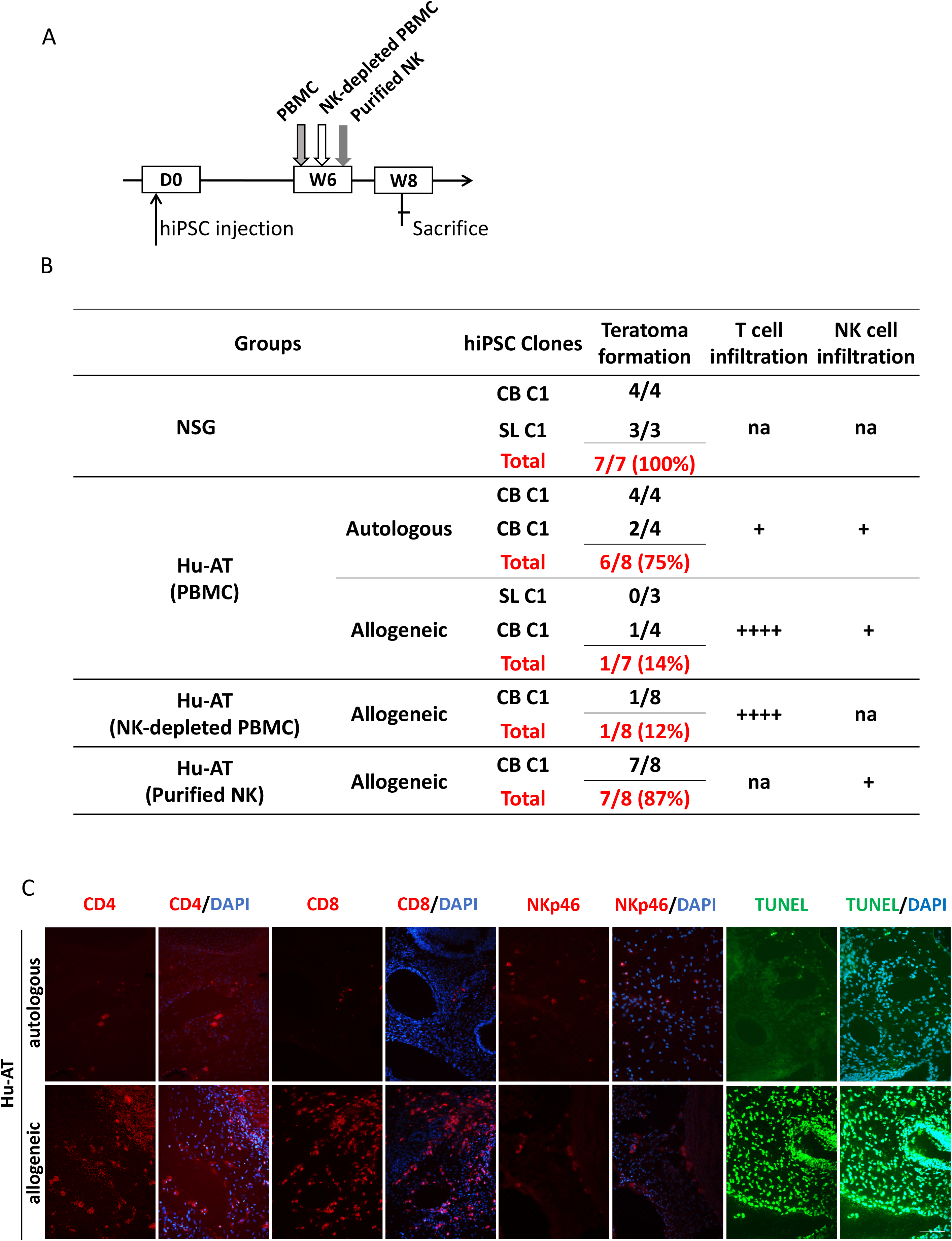
Established teratomas are not the target of NK cells in Hu-AT mice. **(A)** Experimental design and timeline of the injection of hiPSCs in Hu-AT mice. Approximatively 1×10^6^ hiPSCs were first injected under the renal capsule of NSG mice. 6 weeks later (W6), mice were injected intravenously with PBMCs, NK-depleted PBMCs or purified NK cells. 2 weeks after the adoptive transfer of immune cells (W8), mice were sacrificed and the incidence of teratomas and the infiltration of immune cells determined. **(B)** Incidence of teratoma formation in the renal capsule of Hu-AT mice. Shown is the proportion of teratomas derived from the indicated hiPSC clones and their relative infiltration by T and NK cells. na = not applicable. **(C)** Representative photos showing the infiltration of CD4^+^, CD8^+^ T cells, NKp46^+^ NK cells (in red) and apoptosis (as determined by TUNEL in green) in sections of teratomas collected from Hu-AT mice injected with allogeneic or autologous 1×10^7^ PBMCs. Of note, only one teratoma was retrieved in mice injected with allogeneic PBMCs and used for the staining. DAPI staining was performed to visualize nuclei (in blue). Showed are photos taken at 20X. Scale bar, 100 μm.

## DISCUSSION

The ability to generate hiPSCs at an affordable cost and in a timely manner will be necessary to assure the success of future hiPSC-derived cell therapies. Moreover, it will be essential to demonstrate that hiPSC-based therapies, whether using autologous donors or through the development of universal cell lines, are safe [11; 12; 13]. Here, we provide evidences that hiPSCs are the target of NK cells *in vitro* and in humanized mice. In opposition to what was found using murine iPSCs, human NK cells could prevent the formation of teratomas without prior activation [5]. Other than the species differences, such disparity may be explained by the use of a different injection site in our study (renal capsule versus subcutaneous) and perhaps by the high expression level of NK activating ligands (i.e. MICA/B) in hiPSCs. In absence of NK cells, we found that teratomas from allogeneic but not autologous hiPSCswere rejected by T cells in both Hu-AT and Hu-BLT mice. Our results also demonstrated that inhibition of teratomas formation was possible only if hiPSCs were injected prior to the adoptive transfer of total PBMCs or purified NK cells, consistent with the prior observation that hiPSCs-derived myoblasts are not the target of NK cells *in vitro* and *in vivo* (Benabdallah et al. submitted). Moreover, in the absence of recombinant human IL-15 to support the proliferation of NK in our mice [14], our results also suggest that the relatively low number of NK cells injected during the adoptive transfer procedure was sufficient to prevent the growth of teratomas. These results suggest that if immunosuppressive drugs were to be used to increase engraftment of iPSC-derived cells, it would be very important to insure no inhibition of the NK cell compartment to lower the risk of forming a teratoma.

## ACKNOWLEDGMENTS

We are grateful to the flow cytometry platform and animal facility for providing technical support and to Renée Dicaire for handling clinical samples. This work was supported by a grant from the Canadian Institute of Health Research #MOP-126096 to C.M.B. and E. H. by a grant from le réseau ThéCell du Fonds de la recherche du Québec – Santé (FRQS) and the support from la Fondation Charles Bruneau for access to technological platforms. C.M.B. was supported by a senior scientist award from the FRQS.

## AUTHOR CONTRIBUTIONS

B.B., C.D.L., C.C. and Y.L. performed experiments. B.B., H.E. and C.B. designed the studies, J.V.G. provided cells. B.B. and C.B. wrote the manuscript.

## CONFILCT OF INTEREST STATEMENT

The authors declare no competing interests

## REFERENCES

[1] H. Hentze, P.L. Soong, S.T. Wang, B.W. Phillips, T.C. Putti, and N.R. Dunn, Teratoma formation by human embryonic stem cells: evaluation of essential parameters for future safety studies. Stem Cell Res 2 (2009) 198–210.

[2] M.O. Lee, S.H. Moon, H.C. Jeong, J.Y. Yi, T.H. Lee, S.H. Shim, Y.H. Rhee, S.H. Lee, S.J. Oh, M.Y. Lee, M.J. Han, Y.S. Cho, H.M. Chung, K.S. Kim, and H.J. Cha, Inhibition of pluripotent stem cell-derived teratoma formation by small molecules. Proc Natl Acad Sci U S A 110 (2013) E3281–90.

[3] A. Bedel, F. Beliveau, I. Lamrissi-Garcia, B. Rousseau, I. Moranvillier, B. Rucheton, V. Guyonnet-Duperat, B. Cardinaud, H. de Verneuil, F. Moreau-Gaudry, and S. Dabernat, Preventing Pluripotent Cell Teratoma in Regenerative Medicine Applied to Hematology Disorders. Stem Cells Transl Med 6 (2017) 382–393.

[4] R. Dressel, J. Nolte, L. Elsner, P. Novota, K. Guan, K. Streckfuss-Bomeke, G. Hasenfuss, R. Jaenisch, and W. Engel, Pluripotent stem cells are highly susceptible targets for syngeneic, allogeneic, and xenogeneic natural killer cells. FASEB J 24 (2010) 2164–77.

[5] C. Groschel, D. Hubscher, J. Nolte, S. Monecke, A. Sasse, L. Elsner, W. Paulus, C. Trenkwalder, B. Polic, A. Mansouri, K. Guan, and R. Dressel, Efficient Killing of Murine Pluripotent Stem Cells by Natural Killer (NK) Cells Requires Activation by Cytokines and Partly Depends on the Activating NK Receptor NKG2D. Frontiers in immunology 8 (2017) 870.

[6] V. Kruse, C. Hamann, S. Monecke, L. Cyganek, L. Elsner, D. Hubscher, L. Walter, K. Streckfuss-Bomeke, K. Guan, and R. Dressel, Human Induced Pluripotent Stem Cells Are Targets for Allogeneic and Autologous Natural Killer (NK) Cells and Killing Is Partly Mediated by the Activating NK Receptor DNAM-1. PLoS One 10 (2015) e0125544.

[7] H.C. Tseng, A. Arasteh, A. Paranjpe, A. Teruel, W. Yang, A. Behel, J.A. Alva, G. Walter, C. Head, T.O. Ishikawa, H.R. Herschman, N. Cacalano, A.D. Pyle, N.H. Park, and A. Jewett, Increased lysis of stem cells but not their differentiated cells by natural killer cells; de-differentiation or reprogramming activates NK cells. PLoS One 5 (2010) e11590.

[8] L.D. Shultz, M.A. Brehm, J.V. Garcia-Martinez, and D.L. Greiner, Humanized mice for immune system investigation: progress, promise and challenges. Nat Rev Immunol 12 (2012) 786–98.

[9] J.K. Skelton, A.M. Ortega-Prieto, and M. Dorner, A Hitchhiker’s guide to humanized mice: new pathways to studying viral infections. Immunology 154 (2018) 50–61.

[10] T. Zhao, Z.N. Zhang, P.D. Westenskow, D. Todorova, Z. Hu, T. Lin, Z. Rong, J. Kim, J. He, M. Wang, D.O. Clegg, Y.G. Yang, K. Zhang, M. Friedlander, and Y. Xu, Humanized Mice Reveal Differential Immunogenicity of Cells Derived from Autologous Induced Pluripotent Stem Cells. Cell Stem Cell 17 (2015) 353–9.

[11] Q. Liang, C. Monetti, M.V. Shutova, E.J. Neely, S. Hacibekiroglu, H. Yang, C. Kim, P. Zhang, C. Li, K. Nagy, M. Mileikovsky, I. Gyongy, H.K. Sung, and A. Nagy, Linking a cell-division gene and a suicide gene to define and improve cell therapy safety. Nature 563 (2018) 701–704.

[12] T. Deuse, X. Hu, A. Gravina, D. Wang, G. Tediashvili, C. De, W.O. Thayer, A. Wahl, J.V. Garcia, H. Reichenspurner, M.M. Davis, L.L. Lanier, and S. Schrepfer, Hypoimmunogenic derivatives of induced pluripotent stem cells evade immune rejection in fully immunocompetent allogeneic recipients. Nat Biotechnol 37 (2019) 252–258.

[13] X. Han, M. Wang, S. Duan, P.J. Franco, J.H. Kenty, P. Hedrick, Y. Xia, A. Allen, L.M.R. Ferreira, J.L. Strominger, D.A. Melton, T.B. Meissner, and C.A. Cowan, Generation of hypoimmunogenic human pluripotent stem cells. Proc Natl Acad Sci U S A 116 (2019) 10441–10446.

[14] I. Katano, C. Nishime, R. Ito, T. Kamisako, T. Mizusawa, Y. Ka, T. Ogura, H. Suemizu, Y. Kawakami, M. Ito, and T. Takahashi, Long-term maintenance of peripheral blood derived human NK cells in a novel human IL-15-transgenic NOG mouse. Sci Rep 7 (2017) 17230.

