## Supplementary material for "NK cells prevent the formation of teratomas derived from human induced pluripotent stem cells": supp figures

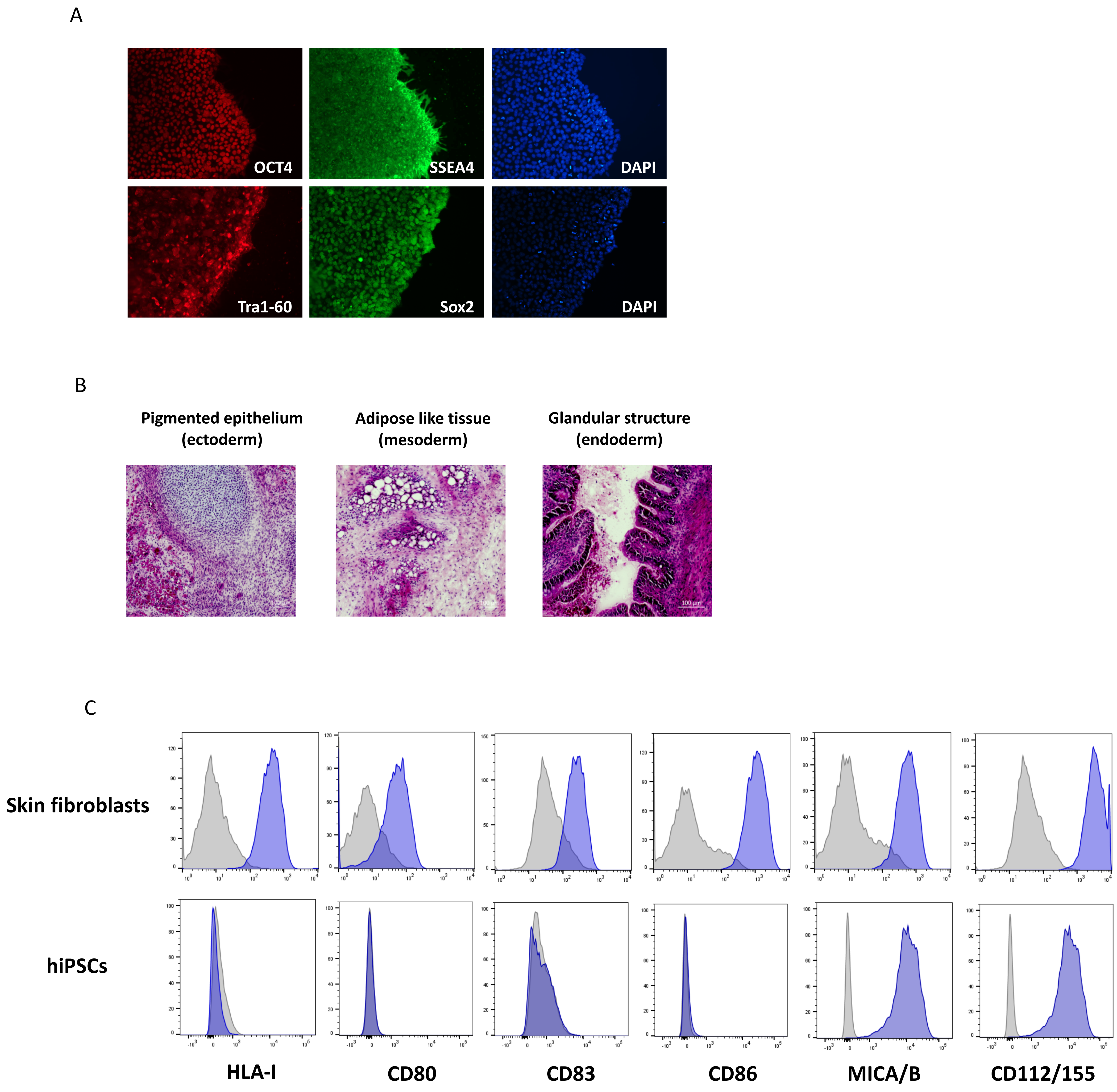


**Figure S1. Phenotypic characterization of hiPSCs.**
(A) Representative photos showing the expression of pluripotency marker (Tra1-60, OCT4 in red and Sox2, SSEA4 in green) in skin fibroblasts-derived hiPSCs. Cells were cultured in feeder-free conditions on Geltrex-coated dishes in E8 medium and passaged every 3-4 days.
(B) Hematoxylin and eosin staining of a teratoma derived from hiPSC clone at passage 17 injected under the renal capsule of a NSG mouse. Representative photo showing tissues from the three embryonic germ layers is shown.
(C) Phenotypic characterisation of human skin fibroblasts and derived hiPSCs. Cells were stained with the indicated mAbs (in blue) or IgG isotype controls (in grey) and analyzed by flow cytometry. Acquisition from one representative experiment is shown.

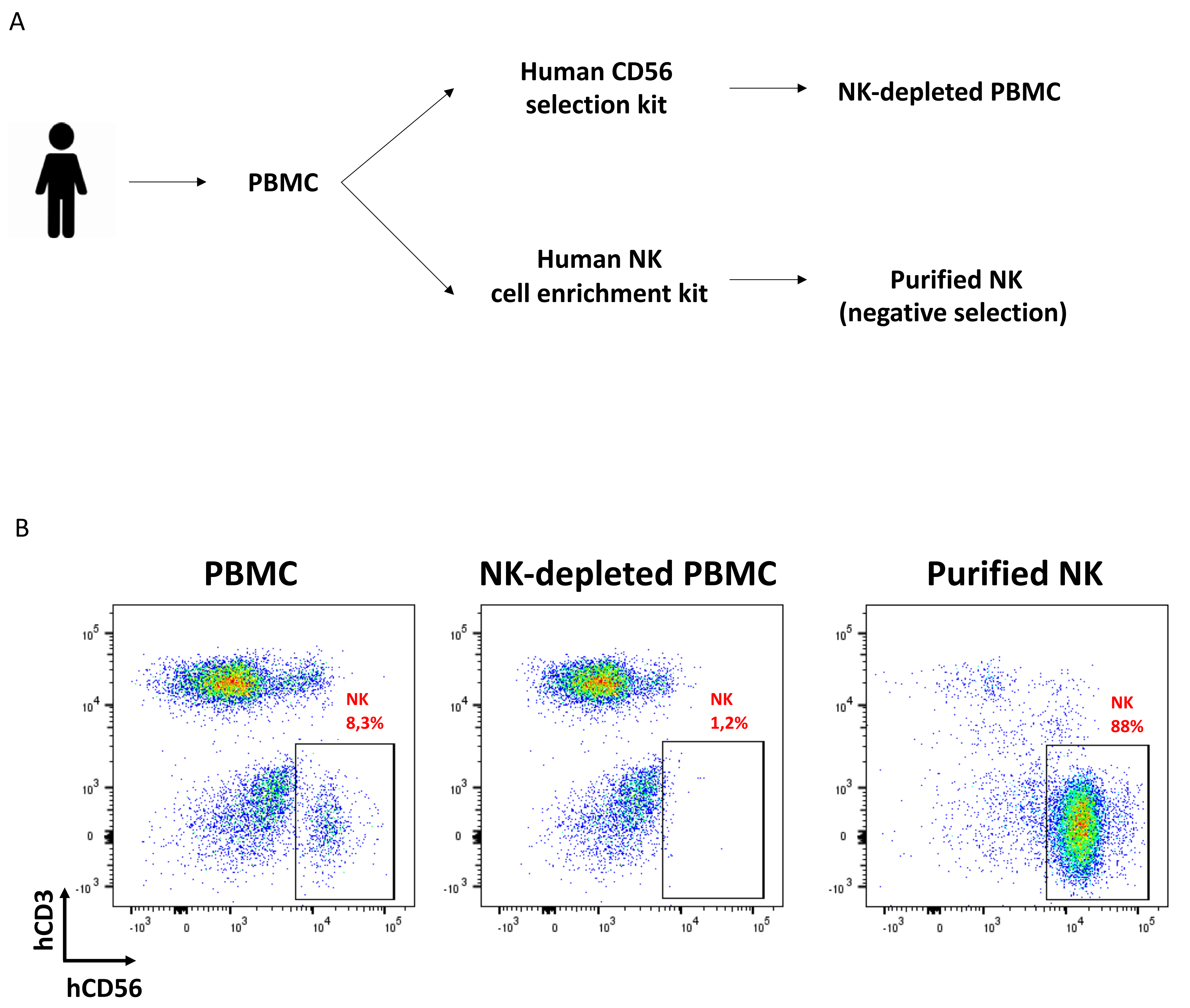


**Figure S2. Purification strategies for the adoptive transfer of human immune cells.**
(A) Schematic illustration of the isolation protocols applied for the magnetic depletion or purification of NK cells from PBMCs used for the reconstitution of the Hu-AT mice.
(B) Representative cell acquisition by flow cytometry of the PBMCs, NK-depleted PBMCs and purified NK showing the proportion of NK cells in each cell population.

A

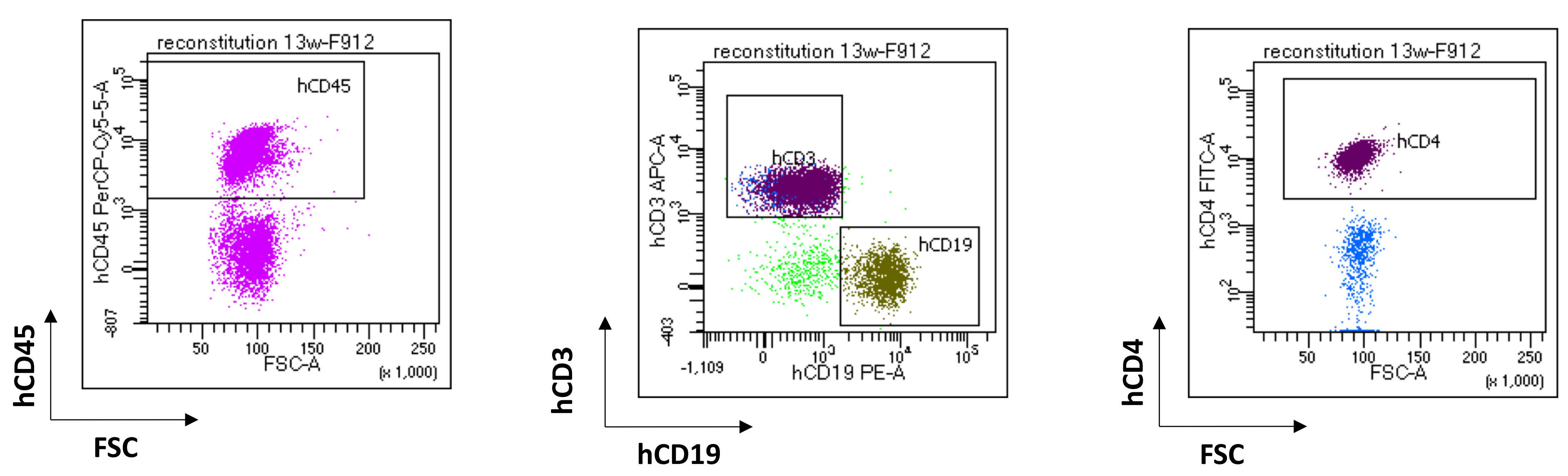


B

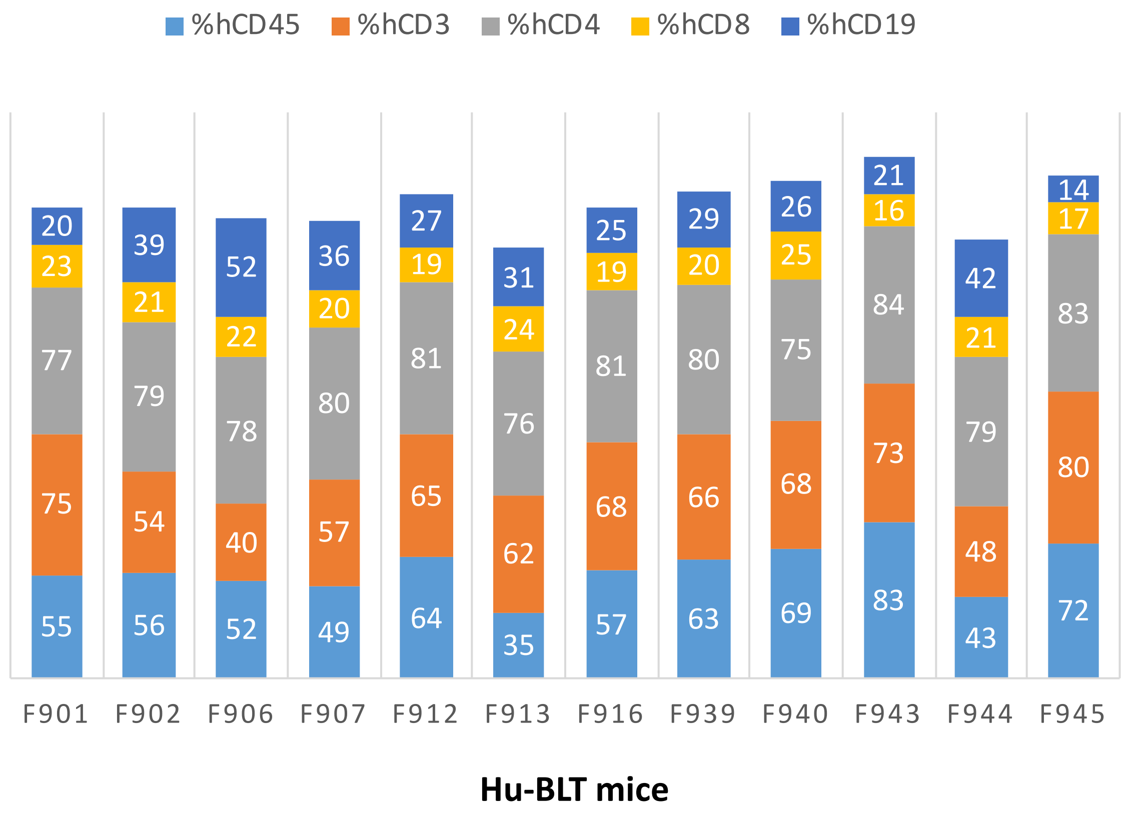


**Figure S3. Immune reconstitution in the peripheral blood of Hu-BLT mice.**
(A) Immune reconstitution of a Hu-BLT mouse 13 weeks following the intravenous injection of fetal liver CD34$^{+}$ cells and the transplantation of autologous thymic tissue under the renal capsule. Representative plots showing human T cells (CD3, CD4) and B cells (CD19) reconstitution in peripheral blood are shown. (B) Frequencies of the major leucocytes subsets found in the peripheral blood of representative Hu-BLT mice 13 weeks following their reconstitution as determined by flow cytometry.

A

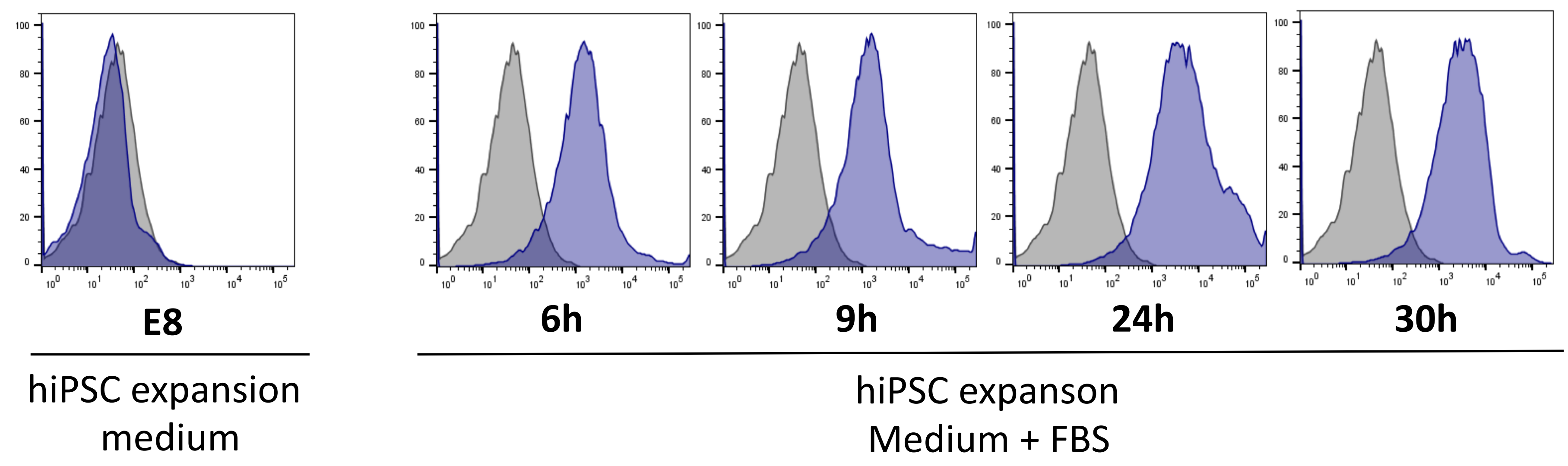


**Figure S4. Increased expression of HLA-1 in differentiated hiPSCs.**
(A) Histograms showing the rapid expression of HLA-I molecules at different time following the addition of 10% fetal bovine serum compared to control cells maintained in E8 expansion medium as determined by flow cytometry. Cells were stained with the anti-HLA-I mAb (in blue) or IgG isotype control (in grey).
